# Branch-Enabling N-Methyltransferase in Golgi Reconciles Divergent Models of Galanthamine Biosynthesis

**DOI:** 10.64898/2025.12.20.695673

**Authors:** Basanta Lamichhane, Archana Niraula, Natacha Merindol, Sarah-Eve Gélinas, Patrick Lagüe, Simon Ricard, Hugo Germain, Isabel Desgagné-Penix

## Abstract

Galanthamine, a therapeutic Amaryllidaceae alkaloid produced exclusively by species within the *Amaryllidoideae* subfamily, is a key treatment for early-stage symptoms of Alzheimer’s disease. Elucidating its biosynthetic pathway is essential for strategies aimed at enhancing production through metabolic engineering. Galantamine derives from the metabolic precursor 4′-*O*-methylnorbelladine, which undergoes cytochrome P450-mediated para-ortho’ C-C phenol coupling to yield nornarwedine. Two competing terminal routes have been proposed: (i) reduction of nornarwedine to norgalanthamine, followed by *N*-methylation, or (ii) *N*-methylation of nornarwedine to narwedine prior to reduction. Here, we identify three *N*-methyltransferase (NMT) candidates from *Leucojum aestivum*: *La*NMT, related to coclaurine NMTs, and two γ-tocopherol methyltransferases (TMT) homologs, *La*TMT1 and *La*TMT2. Subcellular localization studies revealed distinct compartmentalization, with *La*NMT targeted to the ER-cytosol, *La*TMT1 to plastids, and *La*TMT2 to the Golgi apparatus. *In vitro* enzyme assays demonstrated that *La*TMT2 methylates both nornarwedine and norgalanthamine, with a kinetic preference for nornarwedine. Agroinfiltration for transient expression in *Nicotiana benthamiana* further confirmed *La*TMT2 as a catalytically efficient and substrate-promiscuous enzyme that supports both terminal routes. These findings identify *La*TMT2 as a key branch-enabling *N*-methyltransferase that reconcile long-standing models of galanthamine biosynthesis and provides a strategic target for metabolic engineering strategies to enhance galanthamine production.

**Significant statement:** This study identifies *La*TMT2, a Golgi-localized γ-tocopherol methyltransferase homolog, as a branch-enabling N-methyltransferase that resolves competing models of galanthamine biosynthesis. By revealing an unanticipated Golgi-associated step in Amaryllidaceae alkaloid metabolism, it redefines the subcellular organization of specialized metabolic pathways and provides a strategic enzymatic target for metabolic engineering of high-value therapeutic alkaloids.

## 1 Introduction

Amaryllidaceae alkaloids (AAs) are specialized isoquinoline-derived metabolites produced predominantly within the Amaryllidoideae subfamily^1^. AAs exhibit diverse biological activities^2^, including antitumor, antiviral, antibacterial, antifungal, antimalarial, analgesic, acetylcholinesterase inhibitory and cytotoxic effects^3–7^. Among AAs, galanthamine is of particular importance as an FDA-approved therapeutic used for symptomatic management of early-stage Alzheimer’s disease. Like other alkaloids, the structural and pharmacological diversity of AAs is shaped by tailoring enzymes such as methyltransferases (MTs), which modulate a molecule’s stereo-electronic profile, encourage or inhibit specific conformations, enhance steric bulk, increase overall hydrophobicity, and invert the polarity of an electronegative moiety^8^. Despite galanthamine’s importance, the enzymatic step responsible for efficient terminal *N*-methylation has remained ambiguous^9,10^, highlighting a key outstanding question in AA biosynthesis.

Plant MTs are classified based on the atom they methylate (*e.g.,* O, N, C, or S), and typically use *S*-adenosyl-L-methionine (SAM) as a methyl donor^11^. Within plant specialized metabolism, *O*- and *N*-MTs are particularly prominent, catalyzing essential transformations in diverse alkaloid pathways, including tropane, monoterpenoid indole, purine, isoquinoline and benzylisoquinoline alkaloids (BIAs). For example, in tropane alkaloid biosynthesis, a putrescine *N*-MT catalyzes the initial *N*-methylation, ultimately leading to tropinone, which then branches into the tropine and pseudotropine lineages^12,13,14^. Similarly, in monoterpenoid indole alkaloids (MIA), *N*-MTs evolved from γ-tocopherol MTs (TMTs) catalyze the methylation of intermediates such as ajmaline and ervincine^15–17^. For example, in *Catharanthus roseus*, Cr2270, a homolog of the TMT family from *Arabidopsis thaliana,* exhibits high substrate specificity for *N*-methylation in the vindoline pathway^18^. Likewise, TMT homologs, like picrinine-*N*-methytransferase, have also been identified in *Vinca minor* and *Rauvolfia serpentina*, where enzymes such as *Vm*PiNMT and *Rs*PiNMT catalyze the formation of *N*-methylpicrinine^18,19^. *N*-MTs also play crucial roles in purine alkaloid biosynthesis, particularly in the caffeine pathway, where they form a distinct subgroup within the broader MT family more closely related to *C*-MTs^20–23^. In BIAs, which share a common L-tyrosine precursor with AAs, *N*-MTs catalyze multiple key steps. In *Papaver somniferum*, BIA pathways leading to morphinan, protoberberine, and benzophenanthridine alkaloids rely on enzymes such as coclaurine *N*-MT (CNMT)^24^ , reticuline *N*-MT^25^, and tetrahydroprotoberberine *N*-MT^26^. Similar orthologs and functional homologs have been reported across *Ranunculales*, including *Papaver bracteatum, Glaucium flavum, Thalictrum flavum, Coptis japonica, C. chinensis, C. teeta,* and *Eschscholzia californica*^27^.

Structural studies of BIA *N*-MTs provide mechanistic insight into substrate activation and specificity. CNMT from *C. japonica* and pavine *N-*MT (PNMT) from *T. flavum* are canonical SAM-dependent enzymes that share a Rossmann-fold SAM-binding domain and an α-helical substrate-binding region. Despite their homology, CNMT and PNMT display distinct catalytic strategies: CNMT features a compact active site (∼387 Å²) stabilized by conformational rearrangements of helix α4 and adjacent loop regions, whereas PNMT accommodates substrates within a larger pocket (∼1081 Å²) through substrate-induced ordering of flexible loops. Both utilize conserved acidic and histidine residues, specifically the E204/E207/H208 triad in CNMT and the E205/H206 dyad in PNMT, for general base catalysis. Additionally, residues such as Y79/E80 in PNMT and E204 in CNMT may modulate electrostatic stabilization and substrate positioning^28,29^. Together, these features reflect convergent strategies for substrate activation and specificity across structurally homologous *N*-MTs. These residues are conserved across *N*-MTs involved in BIA biosynthesis.

The role of *N*-MTs in AA biosynthesis has been debated, particularly regarding the final *N*-methylation step leading to galanthamine, where two recent studies reached divergent conclusions. The branching of the AA pathway starts from the common precursor 4′-*O*-methylnorbelladine (Figure 1) which undergoes cytochrome P450 enzyme*-*catalyzed *para*-*ortho*′ (*p-o*′) C-C phenol coupling to yield nornarwedine (also known as *N*-demethylnarwedine), which is subsequently reduced by a short-chain dehydrogenase/reductase to norgalanthamine^30^ (Figure 1). However, two alternative routes have been proposed: (i) reduction to norgalanthamine followed by *N*-methylation to yield galanthamine^10^, (ii) *N*-methylation of nornarwedine to narwedine followed by reduction to galanthamine^9^. While our previous work supported the first route with the identification of a CNMT-like enzyme capable of converting norgalanthamine to galanthamine^10^, Mehta *et al.* reported evidence for a TMT acting prior to reduction^9^. Critically, direct enzymatic evidence for *N*-methylation of nornarwedine *in planta* has remained lacking, leaving the order of the terminal reactions unresolved.

**Figure 1.**
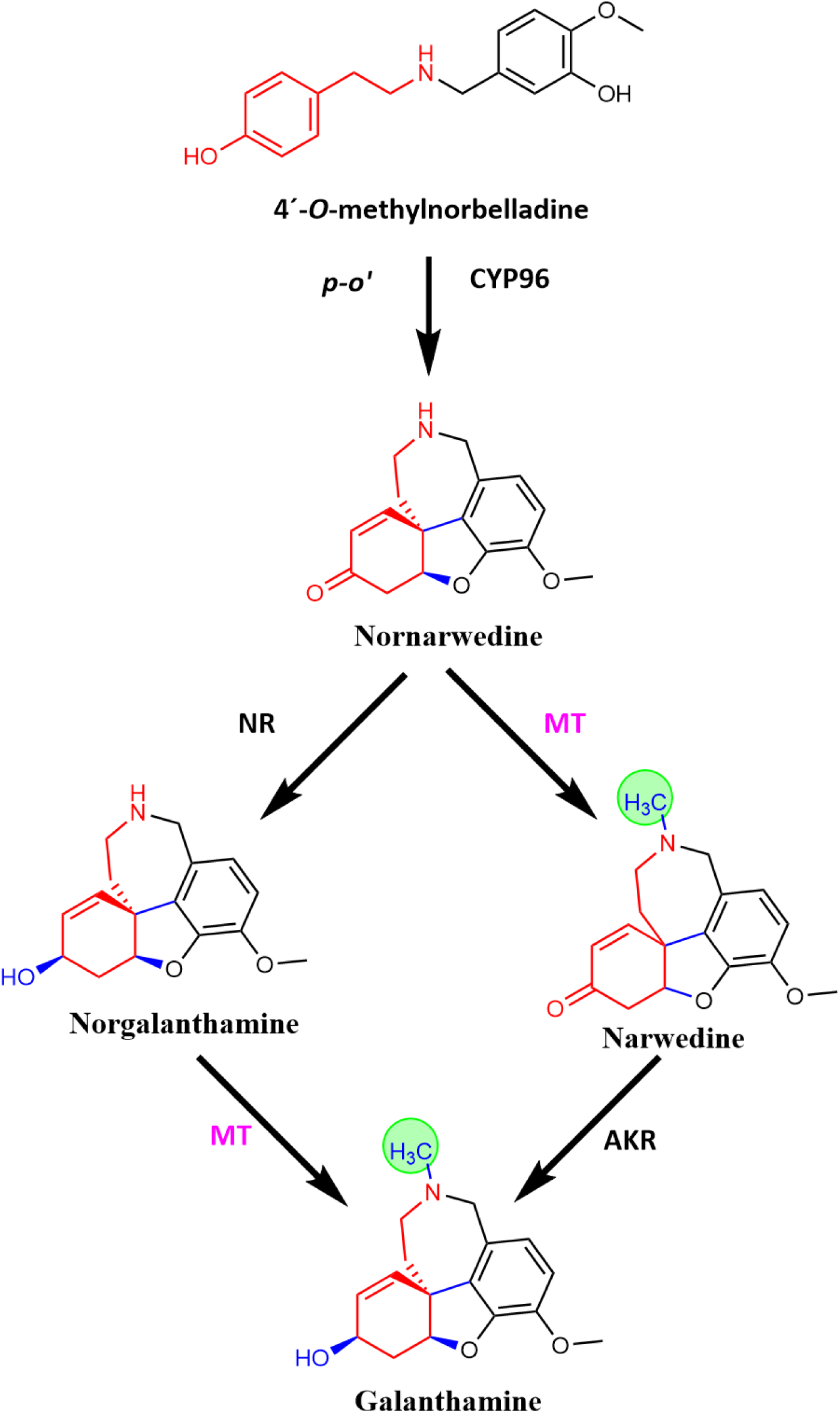
Proposed galanthamine biosynthetic pathway from 4’-*O*-methylnorbelladine. From the cytochrome P450 (CYP) superfamily, the CYP96 family specifically members of the CYP96T subfamily have been reported to catalyze the *para*-*ortho*′ (*p-o’*) oxidative phenol coupling of 4′-*O*-methylnorbelladine to yield nornarwedine. Methyltransferase (MT) in pink that performs methylation in green circle is the enzyme of this study. Structure in red and black is the backbone from Tyramine and 3,4-DHBA respectively and blue is newly formed bonds on action of enzymes. Other abbreviations: Noroxomaritidine/norcraugsodine reductase (NR), aldo-keto reductase (AKR).

To clarify the enzymology underlying the final steps of galanthamine biosynthesis, we systematically compared CNMT-like and TMT candidate enzymes from *Leucojum aestivum* using both nornarwedine and norgalanthamine as substrates. By combining subcellular localization, *in vitro* biochemical assays, and transient expression in *Nicotiana benthamiana*, we identify a Golgi-localized TMT homolog that performs both proposed terminal *N*-methylation reactions, reconciling competing pathway models. These findings resolve a longstanding ambiguity in galanthamine biosynthesis and provide a strategic enzymatic target for metabolic engineering efforts aimed at enhancing galanthamine production.

## 2 Results

### Identification and phylogeny of *N*-methyltransferase candidates

To identify enzymes responsible for the terminal *N*-methylation steps in galanthamine biosynthesis, we mined the *Leucojum aestivum* transcriptome using three TMT reference sequences: *Arabidopsis thaliana* (Uniprot Q9ZSK1), *Catharanthus roseus* Cr2270 (vindoline *N*-methyltransferase; Uniprot W5U2K2), and Cr7756 (picrinine *N*-methyltransferase; Uniprot A0A8X8M501). This search yielded 26, 28, and 29 candidate transcripts per query, respectively. Among them, two *TMT-like* transcripts with the highest identity were selected for further analysis, together with the previously characterized CNMT-like *La*NMT^10^. Phylogenetic reconstruction using ClustalW alignments within a previously established NMT framework^18^ placed *La*TMT1 and *La*TMT2 in the TMT clade, while *La*NMT clustered with BIA-related CNMTs (Figure 2a). Notably, *La*TMT2 subclustered with a previously proposed *N*-methylating TMT candidate from *Narcissus* cv. Tête-à-Tête^9,18^. Coding sequence of *LaTMT1*, *LaTMT2*, and *LaNMT* were 1044, 876, and 1065 bp, encoding 348, 292, and 355 amino acids, respectively. Conserved domain analyses (InterProScan, the NCBI conserved domain database, and ScanProsite/Expasy) confirmed their assignement to the SAM-dependent methyltransferase superfamily. Multiple sequence alignment revealed that all three candidates retained the canonical Rossmann-fold motifs II, III, and IV, and conserved catalytic residues Glu204, Glu207, and His208 (Figure 2b). AlphaFold3 structural predictions further revealed that *La*TMT1 and *La*TMT2 were structurally similar (Root mean square deviation (RMSD) = 1.273 Å), while *La*NMT diverged more substantially (RMSD = 3.485 and 3.067 Å, respectively). Despite divergence, all three candidates exhibited conserved SAM-binding pockets and active site geometry compatible with isoquinoline-like alkaloids (Figure S1). As expected, *La*NMT superposed well with *C. japonica* CNMT (RMSD = 0.498 Å), while *La*TMT1 and *La*TMT2 were more distant (RMSD = 2.839Å; 2.895 Å, respectively) (Figure S1). The N-terminal regions were especially divergent between *La*TMTs and *La*NMT. The folding of SAM and the ligand binding pockets were, however, similar, following the class I methyltransferase α/β Rossmann-fold, with an active site composed mainly of α-helices, and four of the seven conserved β-sheets pointing towards SAM (Figure S1). The catalytic residues Glu204, Glu207, and His208, as well as active site Tyr81 (Tyr72 for *La*TMT1, Tyr17 for *La*TMT2), shared the same orientation, pointing towards superposed CNMT substrate nitrogen.

**Figure 2.**
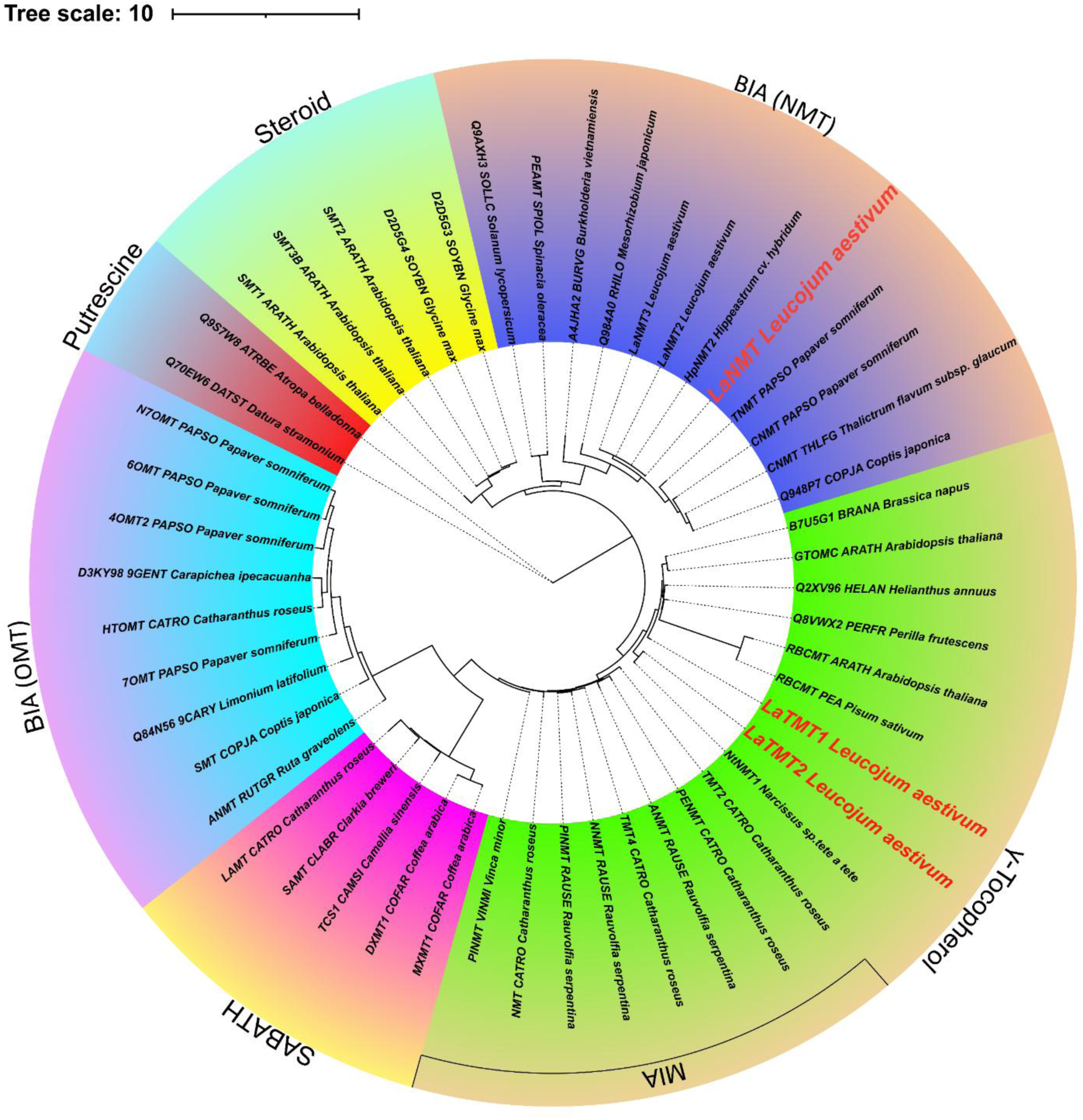
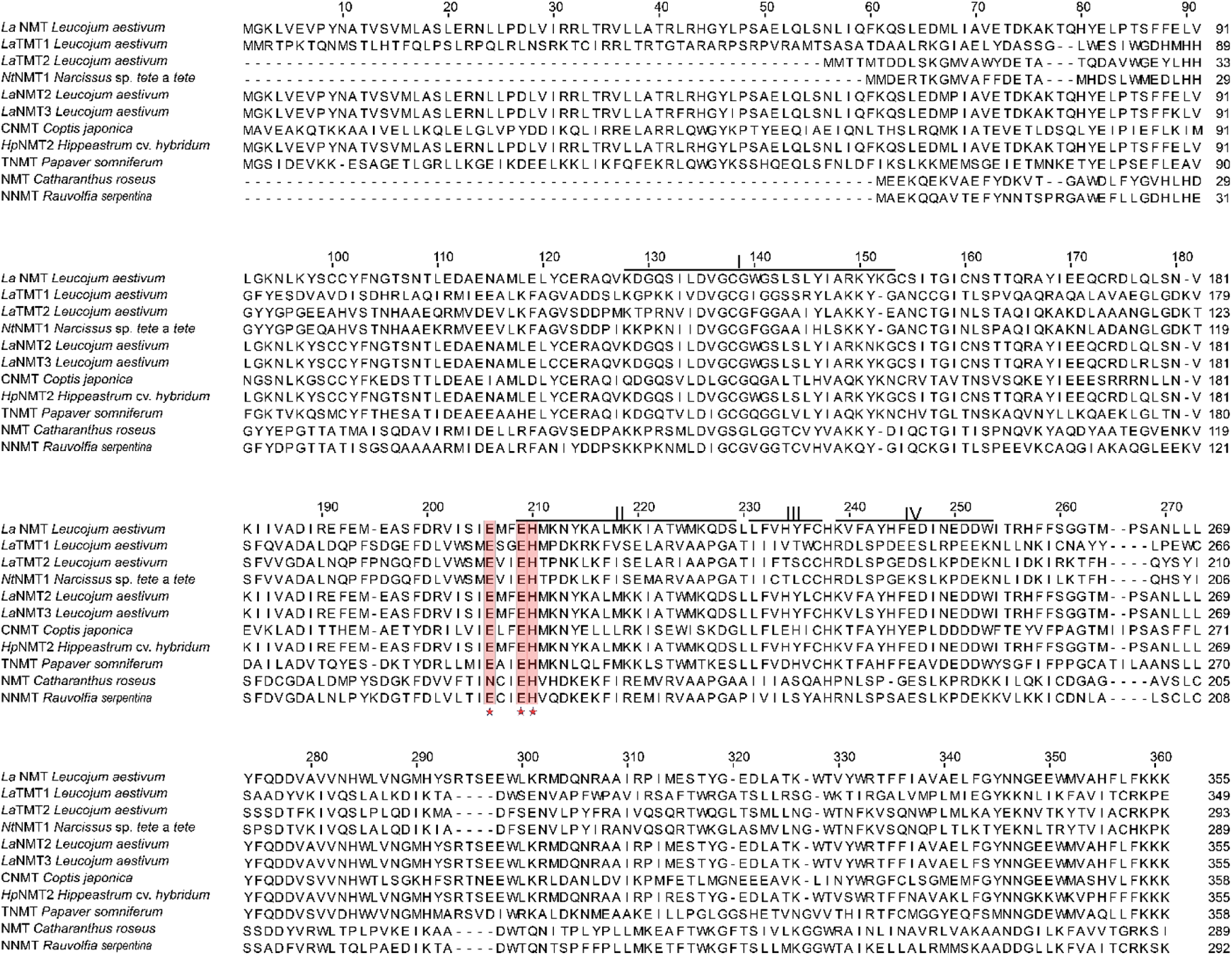
Phylogenetic analysis and sequence alignment of *N*-methyltransferase candidates in Amaryllidaceae alkaloid biosynthesis. (a) The Phylogenetic tree was constructed using the Maximum Likelihood method with 1000 bootstraps. This analysis involved 50 amino acid sequences. Evolutionary analyses were conducted in IQ-Tree and viewed in ITOL software. The transcripts highlighted in bold red font are those included in the present study. The accessions or sequences are listed in Table S1. BIA and MIA stand for benzylisoquinoline alkaloids and monoterpene indole alkaloids. (b) Multiple sequence alignments of 11 sequences, including the first three candidates selected in this study. I, II, III, and IV are the SAM binding sites from different species, and the star represents the residues of the catalytic triad for substrate binding.

Overall, the predicted structures highlight divergence in the N-terminal region among the candidate enzymes, together with a conserved Rossmann fold associated with SAM binding and an active site compatible with isoquinoline-like alkaloids as substrates for SAM-dependent *N*-methylation. Despite the lack of prior mechanistic studies on plant TMTs, the conservation of the sequence and conformation of the putative catalytic residues’ Glu-Glu-His motif in NMTs suggests a possible shared mechanism.

### Leaf-specific AA biosynthesis in *L. aestivum*

To assess the biosynthetic capacity of *L. aestivum* leaves, targeted LC-MS/MS metabolite profiling combined detected *N*-methylated alkaloids lycoramine and galanthamine, along with AA intermediates, including tyramine, phenylalanine, haemanthamine, norgalanthamine, dopamine, vittatine (or crinine), 4’-*O*-methylnorbelladine, L-tyrosine, 3,4-dihydroxybenzoic acid, 4-hydroxybenzoic acid, and norbelladine (Figure S2a-m). Notably, several metabolites predicted in the pathway, including 3,4-dihydroxybenzaldehyde, 3′,4′-*O*-dimethylnorbelladine, 3′-*O*-methylnorbelladine, 9-*O*-demethylhomolycorine, and tazettine, previously reported^31,32^ in the genus, were not detected in any replicate of this organ. A broader absence was noted for additional biosynthetically or structurally related phenolic intermediates such as caffeic acid, *p*-coumaric acid, ferulic acid, isoferulic acid, vanillin, and levodopa, suggesting either abundance below detection limits, tissue-specific biosynthesis, or pathway divergence. 11-Hydroxyvittatine, lycorine, and sanguinine were detected with low-intensity signals in all samples; however, their confirmation was not possible due to insufficient spectral quality. Untargeted metabolite screening was also performed using GC-MS, resulting in the detection of four previously unreported masses (m/z 207, 420, 476, and 490) in the nonpolar fraction (Figure S3a- g). A complete list of both detected and undetected metabolites is provided in Table S2. Altogether, these results confirmed leaves as active sites of galanthamine biosynthesis and provided a metabolic context for enzyme function.

### *La*TMT1, *La*TMT2, and *La*NMT localize in distinct compartments

To determine the subcellular localization of *La*TMT1 and *La*TMT2, each gene was cloned into the pB7FWG2 expression vector to generate C-terminal fusions with EGFP. The resulting constructs were transiently co-expressed in *Nicotiana benthamiana* leaves alongside organelle-specific fluorescent markers^33^: free RFP (nucleus; cytosol), ER-rk *CD3-959* (endoplasmic reticulum-mCherry), and G-rk *CD3-967* (Golgi bodies-mCherry). Confocal laser scanning microscopy was performed 48 hours post-infiltration to assess the subcellular distribution of the fusion proteins. Distinct localization patterns were observed for each of the candidates. The *La*TMT1-EGFP fusion protein displayed a punctate and reticulated fluorescence pattern consistent with chloroplast localization, showing clear overlap with the intrinsic chlorophyll autofluorescence (Figure 3a). In contrast, *La*TMT2-EGFP signal co-localized with Golgi-mCherry, indicating Golgi targeting (Figure 3b). As previously reported, *La*NMT-EGFP exhibited a dispersed fluorescence signal throughout the cytosol and a partial overlap with ER-mCherry (Figure 3c & 3d). This differential compartmentalization suggests distinct functional niches, potentially minimizing cross-reactions while supporting pathway efficiency in distinct metabolic landscape inside the cell.

**Figure 3.**
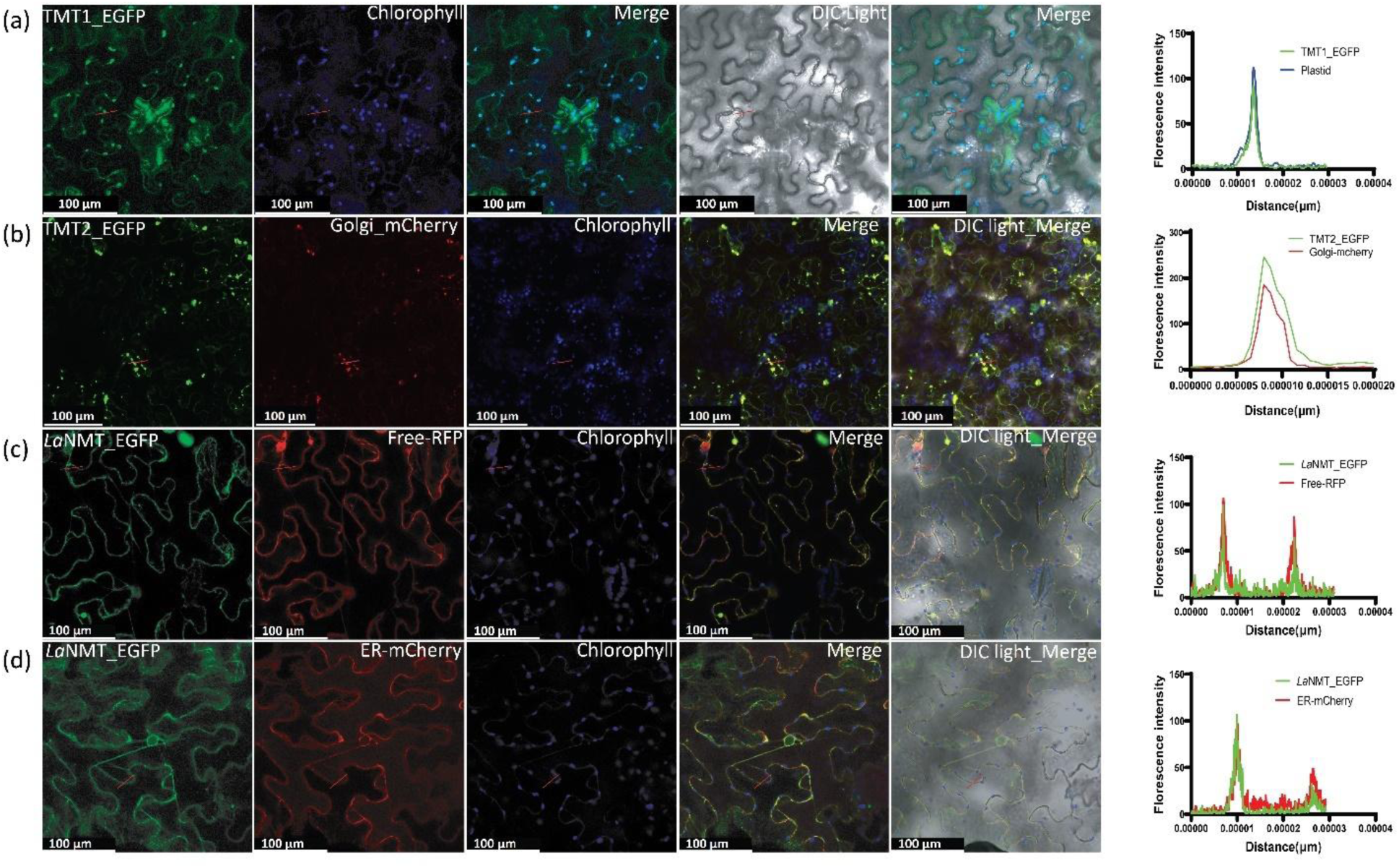
Subcellular localization of *La*TMT1, *La*TMT2 and *La*NMT proteins in *Nicotiana benthamiana*. (a) Panel showing C-terminal-EGFP-fusion *La*TMT1 with chlorophyll autofluorescence. (b) panel showing co-agroinfiltrated C-terminal-EGFP-fusion-*La*TMT2 with Golgi-mcherry marker. (c) Panel showing co-agroinfiltrated C-terminal-EGFP-fusion-*La*NMT with free RFP marker. (d) Panel showing co-agroinfiltrated C-terminal-EGFP-fusion-*La*NMT with ER-mCherry. The histogram of the right in each panel shows the fluorescence intensity profile of respective panel. Scale bars resemble 100 μm. Image processed in Adobe illustrator software.

### *In silico* docking predicts substrate binding

To investigate the catalytic potential of the three candidates, we performed molecular docking using the MOE software. SAM was positioned in the binding pocket based on superimposition with the *Cj*CNMT crystal structure (PDB: 6GKV). Substrate binding poses were analyzed for both norgalanthamine and nornarwedine, the two hypothesized intermediates of galanthamine biosynthesis. The two very similar ligands shared comparable docking scores, positioning, and interaction patterns, reflecting a common binding mode across the tested enzymes. All three enzymes yielded docking scores ranging from -6.3 to -6.8 kcal/mol, with substrate-SAM distances between 3.2 and 3.5 Å (Table 1, Figure 4). This range is compatible with methyl transfer, suggesting all candidates are structurally capable of catalyzing *N*-methylation. In *La*NMT, norgalanthamine binding was stabilized by a conserved hydrophobic core involving Phe234 and Phe257, as well as hydrogen bonds with Tyr98, Glu204, and His208. Notably, Glu204 and Tyr98 formed a direct bond with the substrate amine. Interactions with His208 and Glu204 were lost upon docking nornarwedine. *La*TMT1 featured hydrophobic contacts from Leu71, Trp83, and Tyr260, with Tyr72 and Glu206 forming hydrogen bonds with the substrate (norgalanthamine and nornarwedine) nitrogen. For *La*TMT2, norgalanthamine’s hydrophobic interacting residues included Trp27, Leu31, and Phe204. Tyr17, His33, and His151 contributed polar contacts, with Tyr17 and His151 directly interacting with the substrate amine. Nornarwedine was stabilized by two additional hydrophobic bonds with Trp16 and Glu147. Tyr17 interaction with the amine was conserved. These results support the catalytic potential of *La*NMT for norgalanthamine, as docking led to interactions with the catalytic Glu204 and His208 (Figure 4). *La*TMT1 could accommodate both substrates, although the distance between the amine and SAM was the highest among the three candidates. *La*TMT2 exhibited the most compact and enclosed binding cavity with a favorable substrate-SAM geometry. The ability of all three enzymes to stably bind both norgalanthamine and nornarwedine *in silico* suggests functional promiscuity.

**Figure 4.**
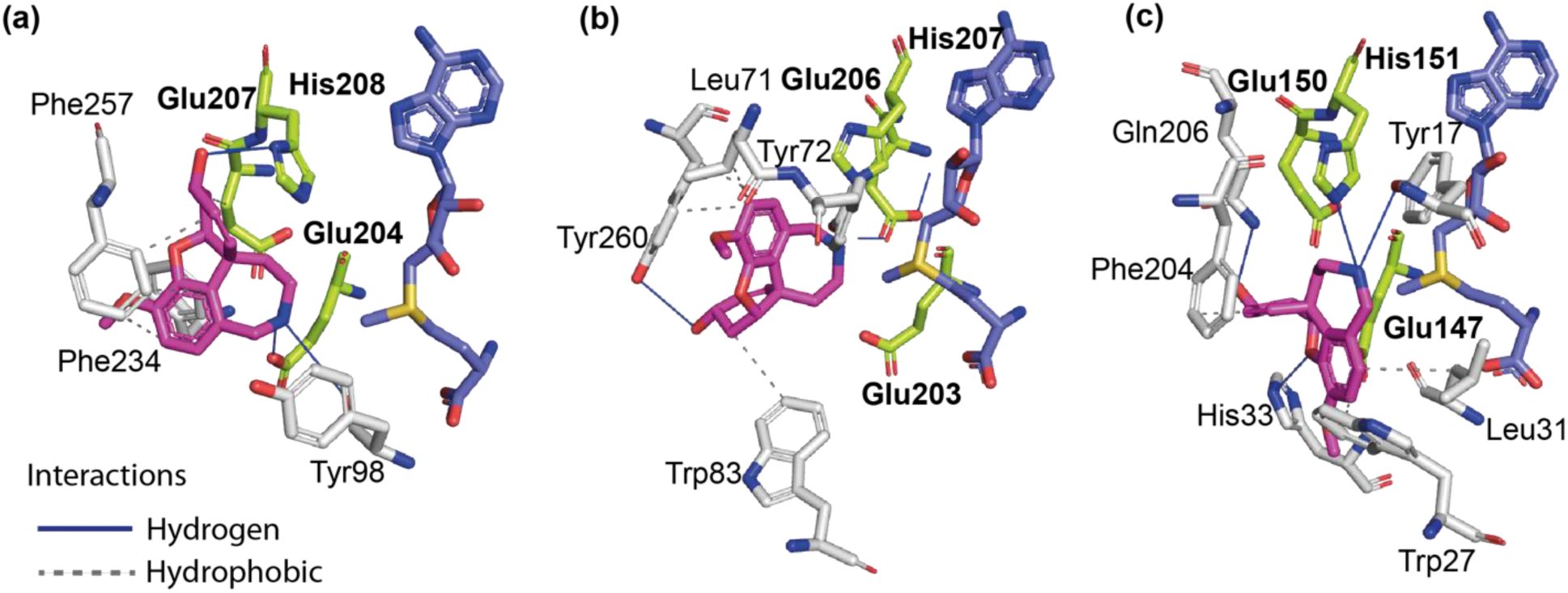
Docking poses and active site interactions of norgalanthamine within the predicted structures of *N*-methyltransferase candidates. (a) *La*NMT, (b) *La*TMT1, and (c) *La*TMT2. In all panels, the ligand norgalanthamine is shown as pink sticks, SAM as purple sticks, and interacting residues are represented as grey sticks. Putative catalytic residues corresponding to the conserved triad are shown as lime sticks. Hydrophobic interactions are depicted as dashed grey lines, and hydrogen bonds as solid blue lines. The orientation is preserved across panels to enable comparison of ligand positioning and interaction networks within the active sites.

**Table 1.** *L. aestivum N*-MT candidates’ interactions with docked norgalanthamine and nornarwedine.

| Substrate (Ligand) | Enzyme | Score and distance | Interactions |  |
| --- | --- | --- | --- | --- |
|  |  |  | Hydrophobic | H-bonds |
| Norgalanthamine | *La*NMT | -6.3<br>(3.3 Å) | Glu207, Phe234, Phe257<br>(n=3) | Glu204 (N), Tyr98 (N), His208 |
|  | *La*TMT1 | -6.8<br>(3.5 Å) | Tyr260, Trp83, Leu71 | Tyr72 (N), Glu206 (N), Tyr260 |
|  | *La*TMT2 | -6.7<br>(3.2 Å) | Trp27, Leu31, Phe204 | Tyr17 (N), His33, His151 (N) |
| Nornarwedine | *La*NMT | -6.3<br>(3.3 Å) | Glu207, Phe234, Phe257<br>(n=3), Phe337 | Tyr98 (N) |
|  | *La*TMT1 | -6.6<br>(3.4 Å) | Leu71, Trp83, Tyr260<br>(n=2) | Tyr72 (N), Glu206 (N) |
|  | *La*TMT2 | -6.5<br>(3.2 Å) | Trp16, Trp27, Leu31, Glu147, Phe204 | Tyr17 (N), His33 |

The binding score is expressed in (kCal/mol). The distance between the substrate and methyl donor was measured between the amine of norgalanthamine or nornarwedine and the carbon of the methyl group of SAM.

### *La*TMT2 exhibits superior catalytic activity *in vitro* and *in vivo*

To evaluate the enzymatic activity of *La*NMT, *La*TMT1, and *La*TMT2, we conducted both *in vitro* and *in vivo* assays using nornarwedine and norgalanthamine as substrates. MBP and EGFP tags served as negative controls, and papaverine was included as an internal standard for LC-MS/MS quantification. In the *in vitro* assays, all proteins were expressed as N-terminal MBP fusion constructs. *La*TMT1 (∼82 kDa) was found to be insoluble. To overcome this, the predicted signal peptide was removed, generating a truncated version (Δ*La*TMT1), which could be purified as a soluble protein. SDS-PAGE confirmed expected band sizes of ∼75 kDa (Δ*La*TMT1), ∼76 kDa (*La*TMT2), and ∼84 kDa (*La*NMT) (Figure S3a). When using nornarwedine as a substrate, LC-MS/MS analysis showed that crude *La*TMT1 and purified Δ*La*TMT1, along with *La*NMT, exhibited detectable activity above MBP background, though not statistically significant (*La*TMT1 0.0022 ± 0.0016; Δ*La*TMT1= 0.0225 ±0.0025; *La*NMT = 0.0203 ± 0.0026, respectively). By contrast, *La*TMT2 catalyzed the formation of narwedine with a 17-fold increase over MBP background and a 9-fold increase over *La*NMT (*p* < 0.0001, Figure 5a), indicating higher activity. Similarly, using norgalanthamine as a substrate, galanthamine production was detectable in reactions with Δ*La*TMT1 and *La*NMT, but *La*TMT2 again showed the highest catalytic efficiency, producing ∼6-fold higher than other enzymes (*p* < 0.0001, Figure 5b).

**Figure 5.**
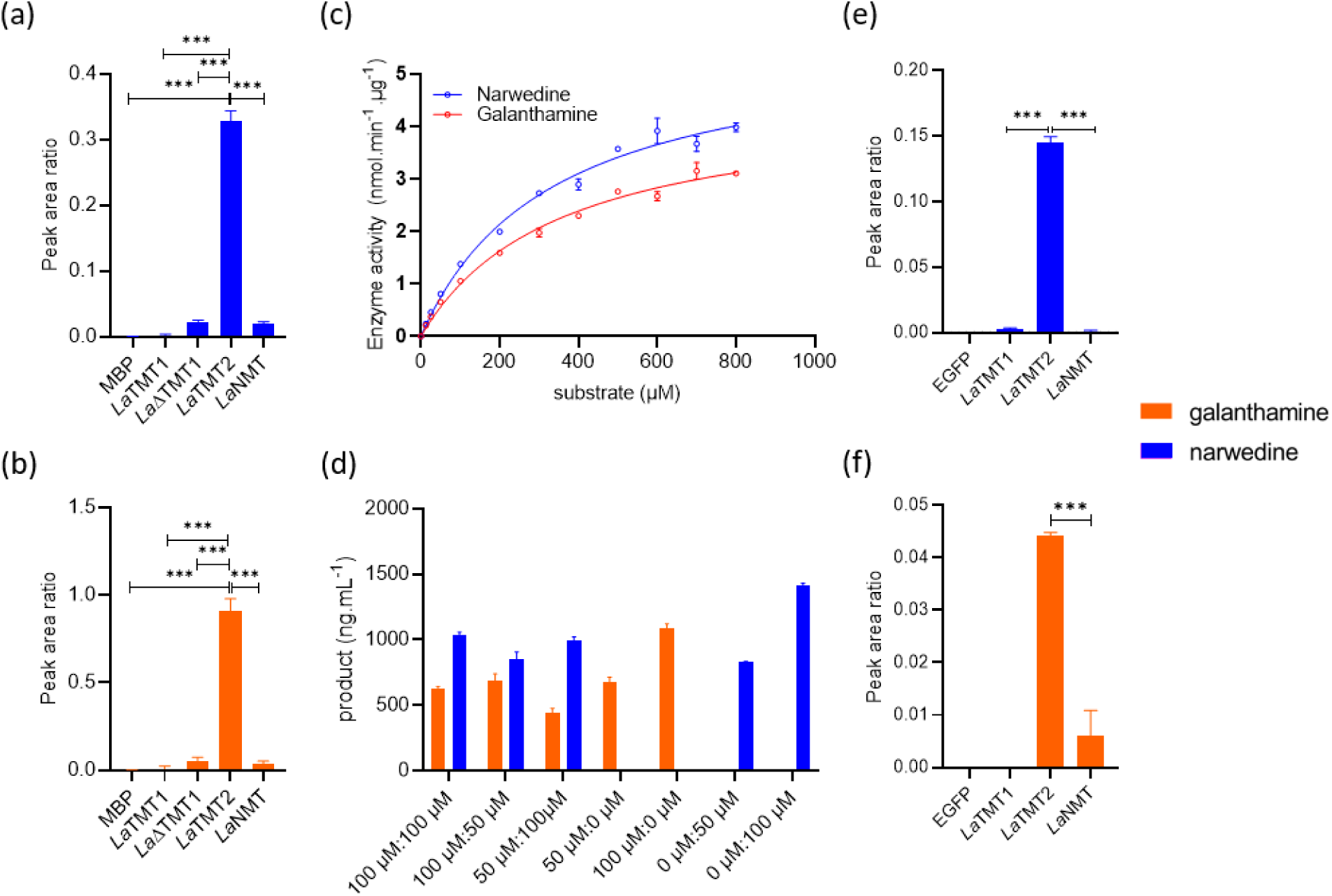
*In vitro* and *in vivo* enzymatic assay by candidate N-methyltransferases. (a) *In vitro* enzymatic assay using nornarwedine as substrate. (b) *In vitro* enzymatic assay using norgalanthamine as substrate. (c) *In vitro* enzyme activity of *La*TMT2 using norgalanthamine and nornarwedine as substrate. (d) Catalytic conversion of substrates norgalanthamine: nornarwedine by *La*TMT2 in a single pot reaction at various ratios. (e) *In vivo* enzymatic assay using nornarwedine as substrate. (f) *In vivo* enzymatic assay using norgalanthamine as substrate.

We further proceed on fully characterizing the *La*TMT2 enzyme by testing its activity on norgalanthamine and nornarwedine at various enzyme concentrations, temperature, incubation time, pH of buffer, and substrate concentrations. At 30°C to 35°C, the enzyme activity for both substrates was optimal with an incubation time of at least 3 hours, reaching a plateau at pH 7.5 (Figure S4a-f). Enzyme kinetic characterization of *La*TMT2 was performed with norgalanthamine and nornarwedine as substrates under optimized *in vitro* conditions. Michaelis–Menten analysis revealed comparable substrate affinities, with Km values of 353.2 ± 28.5 µM for norgalanthamine and 332.2 ± 28.7 µM for nornarwedine. The corresponding maximal velocities (Vmax) were 4.5 and 5.5 nmol·min⁻¹·µg⁻¹, respectively. Turnover numbers (kcat) were 0.225 ± 0.008 s⁻¹ for norgalanthamine and 0.284 ± 0.010 s⁻¹ for nornarwedine (Table 2, Figure 5c). We also examined the substrate preference of the enzyme in mixed-substrate assays, where both substrates were supplied at either similar concentrations or one at half concentration within a single-pot reaction. The amount of product formed revealed more than 1.5% preference for nornarwedine over norgalanthamine (Figure 5d).

**Table 2.**
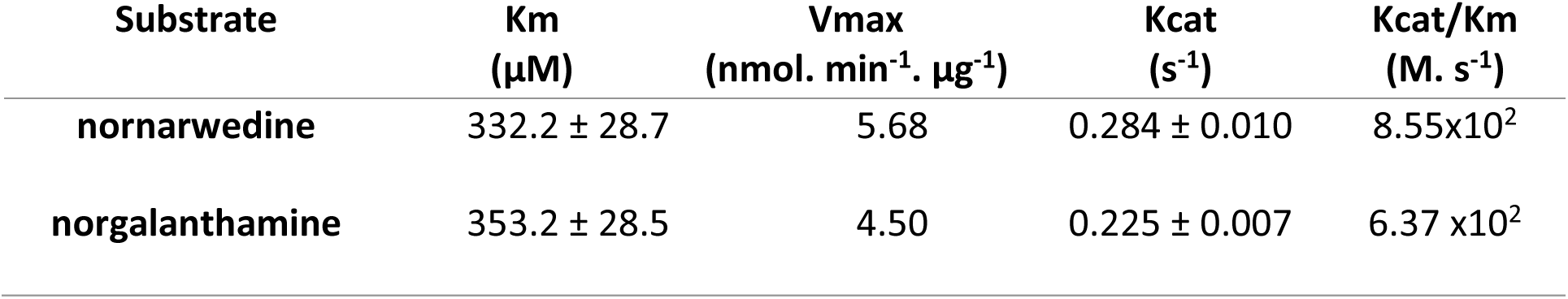
Michaelis-Menten enzyme kinetic parameters for *La*TMT2 with respect to nornarwedine and norgalanthamine.

To confirm these findings *in planta*, the three candidates were transiently expressed in *N. benthamiana*. Immunoblotting confirmed expression of EGFP.3HA (∼30.1 kDa), *La*TMT2-3HA (∼35.7 kDa), *La*TMT1-3HA (∼41.9 kDa), and *La*NMT-3HA (∼44.6 kDa) (Figure S3b). Δ*La*TMT1 was omitted in the *in vivo* assay, as it was not necessary to purify the proteins afterwards. *La*NMT produced galanthamine *in vivo*, but was unable to methylate nornarwedine, whereas *La*TMT*1* catalyzed *N*-methylation of nornarwedine but not its reduced counterpart. In agreement with the *in vitro* results, *La*TMT*2* expression led to the highest accumulation of narwedine, showing a 50-fold and 120-fold increase over *La*TMT1 and *La*NMT, respectively (*p* < 0.0001, Figure 5e). When infiltrated with norgalanthamine, *La*TMT2 again yielded at least a 6-fold higher galanthamine accumulation than all other enzymes (*p* < 0.0001, Figure 5f), confirming its superior activity across both substrates and systems. Taken together, these results establish *La*TMT2 as a robust and substrate-promiscuous *N*-methyltransferase, outperforming both *La*TMT1 and *La*NMT in both *in vitro* and *in planta* settings. Its consistent activity toward two structurally related intermediates highlights its potential as a key enzyme in galanthamine biosynthesis.

## 3 Discussion

MTs constitute a ubiquitous and diverse class of enzymes that are critically involved in a myriad of essential cellular processes across all domains of life. They play fundamental roles in biosynthesis, signal transduction, protein repair, chromatin modulation, and gene silencing^34,35^. Within the Amaryllidoideae, the order of terminal methylation and reduction steps leading to galanthamine has long remained unresolved. We previously reported that the CNMT-like *La*NMT1 catalyzes the *N*-methylation of norgalanthamine to galanthamine *in vitro* and *in vivo*^10^, whereas Mehta *et al.* proposed a TMT-like enzyme acting on nornarwedine *in vivo*^9^. Here, coclaurine- and γ-tocopherol-like MTs from *Leucojum aestivum* to clarify these alternative routes. To clarify these alternatives routes, we identified and functionally compared both coclaurine- and γ-tocopherol-like MTs from *L. aestivum*.

Phylogenetic and structural analyses revealed two *L. aestivum* TMT homologs (*La*TMT1 and *La*TMT2) that diverge evolutionarily from LaNMT. *La*TMT1 and *La*TMT2 grouped closely with neighboring sequences encoding TMT that does *N*-methylation in the MIA pathway like in *C. roseus*^9,18,19,36^. *La*TMT1 grouped with canonical tocopherol MTs from *Pisum sativum, Arabidopsis thaliana,* and *Helianthus annuus*, which are involved in tocopherol (vitamin E) biosynthesis, whereas *La*TMT2 clustered near the TMT from *Narcissus* cv. Tête-à-Tête, previously shown to *N*-methylate narwedine *in vivo*^9^. As expected, *La*NMT diverged significantly and clustered within enzymes of the BIA clade including CNMT from *P. somniferum*^26,37^. Overall, the positioning of the candidates suggests that *La*TMT1 and *La*TMT2 differ from *La*NMT in evolutionary path. Still, the three candidates are homologous to enzymes involved in the *N-*methylation of alkaloids in other plants, indicating that they could contribute to alkaloid diversity in *L. aestivum*.

Structural modeling further supported the functional relevance of all three candidates. Despite divergence in the N-terminal regions, all enzymes retained the conserved Rossmann fold typical of class I MTs, with a well-formed SAM-binding pocket and conserved orientation of key catalytic residues. The CNMT GluxxGluHis catalytic triad and a nearby tyrosine, implicated in substrate positioning, were preserved in both *La*TMTs and *La*NMT, suggesting a possible shared catalytic mechanism despite their phylogenetic separation. Despite the lack of prior mechanistic studies on plant TMTs, the conservation of the putative catalytic residues GluxxGluHis in CNMTs could suggest a functional repurposing of GTMTs for alkaloid *N*-methylation in plant species.

Multiple sequence alignment with functionally characterized reference sequences provided further evidence supporting the identity of the selected transcripts as putative MTs. A defining feature conserved across the MT superfamily is a ∼130 amino acid domain critical for binding the methyl donor SAM^38,39^ and this domain was conserved in all the tree sequences chosen. All three candidates exhibited the conserved GlyxGlyxGly motif, a signature sequence element within the SAM-binding domain, commonly found in both *O*- and *N*-MTs ^40^, further corroborating their functional annotation.

Further metabolite accumulation analysis in leaf tissue shows that *L. aestivum* produces AAs of all three branches: haemanthamine, lycorine, and galanthamine. Previous studies have also reported several metabolites in leaves, as well as in other parts of *L. aestivum*^41,42^. To further complement these metabolite-based insights, we employed molecular docking to explore whether the candidate methyltransferases identified in leaves possess structural features compatible with catalysis of the final *N*-methylation step. Docking simulations suggested that all three MT candidates could be involved in galanthamine biosynthesis. Both norgalanthamine and nornarwedine could be accommodated in the active sites of *La*NMT, *La*TMT1, and *La*TMT2, with similar binding modes and docking scores across the enzymes, suggesting that they could accept structurally related substrates. Notably, *La*TMT2 displayed the shortest distance between the substrate amine and the SAM methyl group, along with a more enclosed binding pocket, indicative of a pre-organized catalytic environment that may favor efficient methyl transfer. In contrast, *La*TMT1 maintained the most extended amine-SAM distances, potentially limiting its catalytic efficiency. *La*NMT, aligning with known BIA NMTs, showed productive interactions with catalytic residues Glu204 and His208, particularly in the presence of norgalanthamine, consistent with our previous study^10^.

Subcellular localization provides key insight into the functional integration of biosynthetic enzymes within specialized metabolic networks. In *P. somniferum* and *Coffea arabica*, their *N*-MTs responsible for alkaloid biosynthesis are cytosolic^43,44^. This cytoplasmic distribution appears to be a common feature among methyltransferases, including alkaloid *O-*MTs such as 4′OMT and 6OMT, which are involved in BIA biosynthesis^30,43^, as well as *O*-MTs involved in the iridoid, vindoline/flavonoid, and anthocyanin pathways^45–47^. Here, the confirmed localization of *La*NMT in both the cytosol and the ER suggests a potential role in intracellular transport processes or interaction with organelle-associated proteins. On the other hand, *La*TMT1 is localized in plastids, like other GTMTs involved in tocopherol biosynthesis^48,49^. The third enzyme, *La*TMT2, is localized to the Golgi complex, a compartment typically associated with protein processing and trafficking, where it may contribute to post-translational modification or sequestration of alkaloid end products. Proteins targeted to the cell wall, vacuole, or other endomembrane compartments are commonly routed through the ER and subsequently trafficked via the Golgi^50^, and several MTs have indeed been identified in the Golgi itself, where they utilize cytosol-imported SAM to methylate substrates^51,52^. Within the broader biosynthetic framework of AAs, these localization patterns indicate a compartmentalized pathway: the early condensation by NBS and reduction by NR occur in the nucleo-cytosolic compartment^53^, cytosolic *O*-methylation generates 4′OMT products, and oxidative coupling by CYP96T1 takes place at the ER^30^, where intermediates likely interface with the cytoplasm and Golgi. In this context, the Golgi-localized *La*TMT2 emerges as a late-acting tailoring enzyme, finalizing *N*-methylation steps in a compartment well suited for product modification, trafficking, and storage. Together, these findings support a model of a highly organized, multi-compartmentalized biosynthetic system in *L. aestivum*, where cytosolic, ER, and Golgi enzymes are spatially integrated to maximize metabolic flux, regulate intermediate pools, and prevent cytotoxic alkaloid accumulation.

*La*TMT1, *La*ΔTMT1 and *La*NMT showed less activity compared to *La*TMT2 to catalyze methylation of both substrates nornarwedine and norgalanthamine *in vitro*, indicating substrate promiscuity. The results are consistent with the proposed catalysis observed following agro-infiltration of an enzyme sharing 80% similarity with *La*TMT2 from *Narcissus* cv. Tête-à-Tête^9^. However, the substrate and activity of this homolog had not been verified. To our knowledge, the current study is the first report to fully characterize the *N*-methyltransferase involved in galanthamine biosynthesis. The similar Km values for both substrates, nornarwedine and norgalanthamine, indicate *La*TMT2 binds both substrates with comparable affinity. The higher turnover number (kcat) for nornarwedine demonstrates that substrate discrimination occurs at the catalytic step rather than during binding. This kinetic profile supports a model in which *La*TMT2 accommodates both terminal intermediates but preferentially drives the GTMT-first route, while still enabling the alternative norgalanthamine branch. This trait is particularly advantageous in the context of metabolic engineering, where enzyme flexibility can facilitate the streamlined reconstruction of pathways in heterologous hosts. Our findings fill the critical knowledge gap in the final steps of galanthamine biosynthesis, uncovering *La*TMT2’s efficiency toward multiple substrates, including norgalanthamine.

Further, *in planta* activity supports the physiological relevance of *La*TMT2’s catalytic function, implying compatibility with the cellular environment and endogenous cofactor pools. The combination of the localization and enzyme activity result suggests the coupling of *La*TMT2-mediated *N*-methylation catalytic function with intracellular trafficking or sequestration of alkaloid end products. Future work should aim to resolve whether *La*TMT2 resides in the cis-, medial-, or trans-Golgi, and to determine whether Golgi-synthesized alkaloids are subsequently transported via vesicular trafficking to other organelles, including the vacuole. *La*TMT2 may occupy a central metabolic node in native alkaloid biosynthesis and possibly evolved for efficiency and flexibility under diverse cellular conditions. Its promiscuous activity profiles observed *in vitro* and *in planta* reinforce the idea that *La*TMT2 is an evolutionarily repurposed γ-tocopherol methyltransferase with efficient and flexible *N*-methylation capabilities relevant to galanthamine biosynthesis.

Our results provide enzyme-level support for the long-hypothesized existence of multiple galanthamine biosynthetic routes, complementing earlier radio- and stable-isotope labeling studies that suggested contributions from both norgalanthamine and nornarwedine^54^. The dual activity of *La*TMT2 implies that *N*-methylation may serve as a metabolic branching point, ensuring pathway robustness under variable precursor availability. Whether a single reductase acts upstream or downstream of both routes, or distinct reductases partition flux, remains an open question and may represent a critical regulatory step. The substrate promiscuity of *La*TMT2 highlights the potential for additional *N-*methylated alkaloid derivatives in Amaryllidaceae, underscoring the value of future substrate scope and structural studies to uncover hidden diversity and enable metabolic engineering. This work positions *La*TMT2 as a valuable tool for synthetic biology efforts aimed at producing high-value alkaloid, galanthamine. The ability of *La*TMT2 to act on both nornarwedine and norgalanthamine also raises intriguing questions about its active site architecture and binding dynamics, offering avenues for future structural and mechanistic studies of enzyme.

## 4 Conclusion

This study resolves the ambiguity surrounding the *N*-methylation step in galanthamine biosynthesis, a pharmacologically important alkaloid produced exclusively by the Amaryllidoideae subfamily of Amaryllidaceae. Through an integrated approach combining phylogenetic, structural, and docking analyses with *in vitro* enzymatic assays and *in vivo* transient expression in *N. benthamiana*, we identified *La*TMT2 as a Golgi-localized TMT homolog with superior catalytic efficiency and substrate promiscuity. *La*TMT2 effectively methylates both nornarwedine and norgalanthamine, reconciling divergent biosynthetic models and defining a flexible terminal step in galanthamine formation. Our study is the first to show its robust *in vitro* function and substrate scope by characterizing it. The consistency of *La*TMT2 activity across expression systems and substrate types highlights its utility as a versatile biocatalyst. Beyond elucidating a missing enzymatic link, this work provides a foundation for rational metabolic engineering aimed at enhancing the sustainable production of galanthamine and related Amaryllidaceae alkaloids.

## 5 Materials and methods

### Identification of candidates and phylogenetic analysis and Multiple Sequence Alignment

To find putative gamma tocopherol-like NMT, G-TMT from *Arabidopsis thaliana* and Cr2270 from *Catharanthus roseus*, and CNMT from *P. somniferum* were used as bait to pull the candidate transcripts from the *Leucojum aestivum* transcriptome database. Sequences were aligned by ClustalW. Phylogenetic analysis was done using IQ-TREE with sequences obtained from UniProt SwissProt or the NCBI database along with published sequences by Liscombe et al.^11,18,55^. The phylogenetic tree was inferred using the Maximum Likelihood method with the JTT matrix-based model. As part of the heuristic search, initial tree topologies were generated automatically using the Neighbor-Joining and BioNJ algorithms, and the best-scoring topology based on log-likelihood was selected for ML optimization. The tree was drawn to scale with branch lengths representing substitutions per site, and visualized using iTOL v6.^56,57^

### Molecular modelling of *La*NMT, *La*TMT1, and *La*TMT2

The established models of CNMT bound to *N*- (6GKZ, 6GKV) and *S*-adenosyl-L-homocysteine from *C. japonica* were used as reference structures ^28^. Folding prediction of *La*NMT, *LaTMT1* and *LaTMT2* candidates was performed using AlphaFold Server Beta powered by AlphaFold3 ^58^. Receptor preparation and docking were completed using MOE 2024.0601 software with the AMBER: EHT10 force field (Chemical Computing Group). *Cj*NMT was used as a reference to position the methyl donor SAM in the active site of the candidate following superimposition. The docking site was predicted using *Site Finder* tools. Ligands (norgalanthamine, nornarwedine) isomeric SMILES codes retrieved from PubChem were submitted to the ZINC20 database to obtain 3D charged data files ^59^. Two protomers predicted at pH = 8.0 were included as possible ligands. *Triangle matcher* was used as a placement method for 200 poses generated with tethered *induced fit* refinement. Ten docking poses for each protomer were compared with the crystalized protein-ligand complex. The pose with the best docking score, coherent with the catalyzed reaction, was selected. PLIP was used to detect the interactions between ligands and receptor residues^60^. Visualization and figure preparation were performed using Pymol 2.3.4 (Schrödinger).

### RNA extraction and cDNA synthesis

RNA was isolated from 80 mg of snap-frozen *Leucojum aestivum* leaves using the RNeasy® Plant Mini Kit (Qiagen, Mississauga, Canada) according to the manufacturer’s protocol. DNAse treatment was performed using Turbo-DNAse according to the standard protocol from the supplier (Invitrogen, TURBO DNA-*free* Kit). cDNA synthesis was carried out using the M-MLV reverse transcriptase Kit (Thermo Fisher Scientific, Canada) according to the manufacturer’s protocol, employing both random hexamers and oligo(dT). The resulting cDNA was stored at – 80°C.

### PCR amplification of key candidate genes *LaNMT*, *LaTMT1*, *LaTMT2*

Coding sequences of *LaNMT*, *LaTMT1*, *LaTMT2* from *L. aestivum* were amplified using primers with attB1 and attB2 overhangs along with a 3HA tag in the C-terminal (Table S3) for *in vivo* enzymatic assays in *N. benthamiana*. For the *in vitro* enzymatic assay, coding sequences were amplified using gene-specific primers with restriction sites overhangs compatible with pMAL-c2x multiple cloning site i.e. EcoRI and SalI for *La*TMT1, EcoRI and XbaI for *La*TMT2 as well as *La*NMT. PCR amplifications were performed using PrimeStar GXL premix (TaKaRa Bio) in a T100 Thermal Cycler (Bio-Rad). Amplifications were verified by running 2 µL of PCR product on a 0.9% agarose gel, with images captured using the Bio-Rad ChemiDoc Imaging System. PCR clean-up and gel purification were carried out following the manufacturer’s instructions using the GeneAid PCR & Gel Kit.

### Cloning and *Agrobacterium* transformation for *in vivo* characterization

PCR-amplified fragments were cloned using the Gateway cloning system according to the manufacturer’s instructions (Thermo Fisher Scientific). In this process, the attP-flanked pDONR221 vector, which contains a kanamycin resistance gene, was used in a BP recombination reaction to produce attL-flanked entry clones with attB-flanked sequences for *LaNMT*, *LaTMT1*, *LaTMT2* and *EGFP*. These entry clones were then transformed into *E. coli* DH5α chemically competent cells, which were cultured on LB agar plates containing 50 µg/mL kanamycin. Positive clones were identified via colony PCR, followed by plasmid extraction using the Presto Mini Plasmid Kit (Geneaid), according to the manufacturer’s protocol. The integrity of each positive entry clone was confirmed by Sanger sequencing (Genome Quebec). After verification, selected plasmids were used in an LR recombination reaction, conducted overnight with LR clonase enzyme, to facilitate the recombination between the attL-flanked entry clone and attR-flanked destination vector (pK7WG2). The resulting LR reaction products were transformed into *E. coli* DH5α cells and selected on LB agar plates containing 50 µg/mL spectinomycin. Positive clones were confirmed through colony PCR using Taq polymerase (Taq’Ozyme, France) and, from positive clones, plasmids were extracted using 10 mL LB broth cultures supplemented with spectinomycin. The confirmed expression clones were then transformed into *Agrobacterium tumefaciens* strain GV3101 by electroporation. For the localization, constructs of *LaNMT*, *LaTMT1* and *LaTMT2* cloned were in pB7FWG2 expression vectors to have C-terminal fusion EGFP protein and were co-transformed with free RFP or ER-mCherry (ER-rk *CD3-959*), or Golgi-mCherry (G-rk *CD3-967*) plasmids into *Agrobacterium tumefaciens* strain GV3101^33^. After 48 hours of incubation on LB plates at 28°C, the presence of the insert was re-confirmed via colony PCR like before.

### Confocal image acquisition

Laser scanning confocal microscopy was performed using a Leica TCS SP8 system (Leica Microsystems) equipped with a HC PL APO CS2 40×/1.30 oil immersion objective. Fluorescence signals were examined in sequential mode from agroinfiltrated leaf discs. GFP and chlorophyll were excited at 488 nm, while RFP/mCherry was excited at 561 nm. Emission was recorded at 520–540 nm for GFP, 580–620 nm for RFP/mCherry, and 680–700 nm for chlorophyll autofluorescence. Subcellular localization was assessed in three independent experiments, and image processing was carried out using LAS AF Lite software (v.4.0.0.11706).

### Cloning, transformation, and protein purification for *in vitro* characterization

PCR-amplified fragments were cloned using the restriction digestion/ligation cloning system according to the manufacturer’s instructions (NEB). The PCR fragments and pMAL-c2X plasmids were double digested with restriction enzymes and ligated by T4 DNA ligase. Transformation and plasmid amplification/purification was performed like before using 100 µg/mL ampicillin in growth media. Plasmids were transformed into *E. coli* Rossetta cells on LB agar plates containing 100 µg/mL ampicillin and chloramphenicol (34 µg/mL). Positive clones were confirmed through colony PCR, and single colony of *E. coli* Rosetta (BL21) harboring the gene of interest were further grown in 5 mL LB broth cultures (seed culture), supplemented with ampicillin and chloramphenicol overnight. After 14 hours, 1 mL of seed culture was transferred in 300 mL LB broth supplemented with antibiotics. As the OD_600_ reached 0.5, 0.25 mM isopropyl-β-D-thiogalactopyranoside, 50 μM zinc chloride, and 50 μM magnesium chloride were added, and the cultures were incubated overnight at 18°C with 210 rpm in a shaking incubator. Cells were harvested and protein extraction and purification was performed^30^. Finally, the pure, concentrated, and desalted proteins were obtained using amicon ultra centrifugal filters (Merk, Millipore) and aliquoted in individual tubes and preserved at –80°C.

### *In vitro* enzymatic assay

Recombinant MTs proteins were expressed as N-terminal maltose-binding protein (MBP) fusion constructs and purified using affinity chromatography as detailed by New England Biolabs, Inc. For *La*TMT1, which remained insoluble, crude protein extracts were used directly at a final concentration of 20 µM. All enzymatic reactions were performed in a total volume of 100 µL under the following standardized conditions: 100 mM sodium phosphate buffer (pH 7.5), 2 mM SAM as the methyl donor, 100 µM of substrate, and 5 µM of purified enzyme. Initially reactions were incubated at 35°C for 2 hours and 30 minutes. Following incubation, reactions were quenched by adding an equal volume of mobile phase acidified to pH 2. Papaverine was added as an internal standard to make final 1 ppm in the reaction mix from a stock of 200 ppm. Samples were centrifuged at 10,000 rpm for 15 minutes to remove precipitated proteins, and the resulting supernatants were subjected to downstream metabolite analysis by LC-MS/MS. To comprehensively characterize the *La*TMT2 enzyme, we performed enzyme kinetics under systematically varied conditions. Serial dilutions of *La*TMT2 (5–0.156 µM, two-fold dilutions) were assayed at 1, 3, and 5 hours to determine optimal enzyme concentration and reaction duration. Temperature dependence was evaluated over a gradient of 20, 30, 40, and 50°C for 1–5 hours. Using the optimal enzyme concentration, reaction time, and temperature, the effect of pH on enzymatic activity was assessed across five buffer conditions (pH 5.0, 5.5, 6.5, 7.5, 8.5, and 9.5). In all assays, S-adenosylmethionine (SAM, 2 mM) served as the methyl donor, and the substrate concentration was maintained at 50 µM. Finally, under the identified optimal conditions (enzyme concentration, reaction time, temperature, and pH), substrate saturation experiments were performed to determine the Michaelis–Menten kinetic parameters, Km and kcat.

### Transient expression of candidate genes in *N. benthamiana* (*in vivo* assay)

Individual *Agrobacterium tumefaciens* GV3101 colonies carrying the desired gene constructs (*LaNMT*, *LaTMT1*, *LaTMT2*, and *EGFP*) were grown in LB broth containing 50 µg/mL spectinomycin, 50 µg/mL rifampicin, and 30 µg/mL gentamicin, and incubated at 28°C for 18-20 hours. Culture was centrifuged at 3000 rpm, the pellet was washed in *Agrobacterium* induction buffer (10 mM MES, pH 5.6, 10 mM MgCl_2_, and 150 µM acetosyringone) and following incubation for 2 hours at room temperature. 3-4 weeks old *N. benthamiana* plants, grown in a controlled environment chamber at 22°C with a 16-hour /8-hour dark cycle, were used. The leaves were infiltrated with *Agrobacterium* suspensions adjusted to an optical density at 600 nm (OD_600_) of 0.6 units, using a needleless syringe to infiltrate the suspensions to the abaxial side of the leaves.

Three leaves from three independently infiltrated plants were used as a triplicate to test the hypothesis. Initially, the *Agrobacterium* suspensions containing the gene constructs were infiltrated into the leaves and incubated for 48 hours, followed by a second infiltration step with 100 µM substrates. After 24 hours, leaves were harvested, snap-frozen, and stored at –80°C for further analysis.

### Western blotting

The frozen samples were ground into a fine powder using a mortar and pestle under liquid nitrogen. The expression of the agroinfiltrated genes in *N. benthamiana* leaves was verified by western blot. For this, 100 mg of tissue was resuspended in 0.5 mL of lysis buffer (100 mM Tris-HCl, pH 7.5, 1 mM phenylmethylsulfonyl fluoride, 10% glycerol, 10 mM β-mercaptoethanol, and 2% polyvinylpolypyrrolidone) and incubated on ice for 30 minutes with gentle inversion. The extracts were then centrifuged at 12,000 g for 30 minutes at 4°C to pellet the debris, and the supernatants were collected. Protein samples were run in an SDS gel with 6% stacking and 10% resolving phase. A specific primary antibody was used to verify protein expression on PVDF membranes after semi-dry transfer, followed by blocking with 5% milk for 2 hours at room temperature. A rabbit anti-HA monoclonal primary antibody raised in rabbit (Cell Signaling) was used overnight at 4°C. Host-specific secondary antibodies conjugated with horseradish peroxidase (HRP), specifically the Immun-Star goat anti-rabbit HRP conjugate from Bio-Rad, were utilized, and the membrane was illuminated with ECL reagent to detect the bands on the membrane.

### Substrates and standards preparation

Narwedine and nornarwedine were synthesized from galanthamine hydrobromide in two steps via a protocol adapted from reported procedures^61,62^.

**Figure 6:**
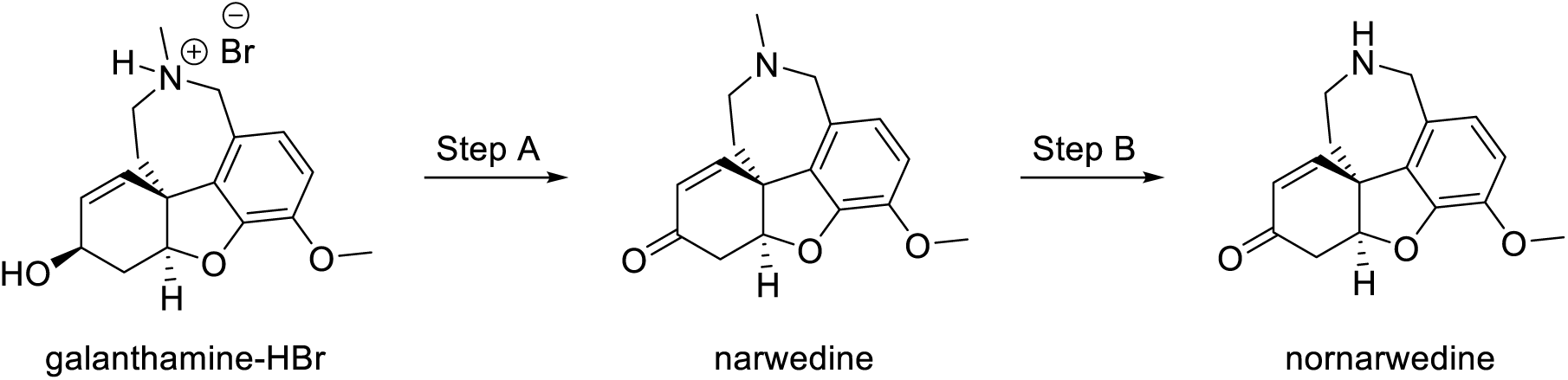
Chemical synthesis of narwedine and nornarwedine.

**Step A.** Galanthamine hydrobromide (200.9 mg, 0.546 mmol) and potassium carbonate (150.7 mg, 1.090 mmol) were added to dry dichloromethane (10 mL), then stirred at room temperature for 1.5 hour. The mixture was filtered over glass wool to remove the remaining solid and the latter was rinsed with dry dichloromethane (3 x 5 mL). The combined dichloromethane solution (25 mL) was brought to 0°C before adding manganese (IV) oxide (94.2 mg, 1.084 mmol), then stirred at 0°C for 5 minutes. The mixture was then stirred at room temperature for 50 hours before adding a second portion of manganese (IV) oxide (47.2 mg, 0.543 mmol). The mixture was stirred at room temperature again for 20 hours before adding a third portion of manganese (IV) oxide (47.7 mg, 0.549 mmol). The solution was finally stirred at room temperature for 30 hours. The reaction mixture was then passed through Celite, rinsed with dichloromethane, and the filtrate was evaporated under reduced pressure to afford narwedine (26.1mg, 17%) without further purification. Spectroscopic data (GCMS, ^1^H-NMR and ^13^C-NMR) matched those reported in the literature^61,62^.

**Step B.** Narwedine (21.0 mg, 0.0736 mmol) and *meta*-chloroperoxybenzoic acid (14.5 mg, 0.0840 mmol) were added to dry dichloromethane (2 mL), then stirred at room temperature for 3.5 hours to form narwedine *N*-oxide *in situ*. To the latter crude solution of narwedine *N*-oxide was added a solution of ferrous sulfate heptahydrate (12.7 mg, 0.0457 mmol) in methanol (1 mL), and the mixture was stirred at room temperature for 30 minutes. A solution of disodium hydrogen phosphate (15.6 mg, 0.110 mmol) in water (1 mL) was added to the reaction, and the mixture was stirred at room temperature for 10 minutes. The precipitate was collected by filtration, then washed with dichloromethane. The pH of the biphasic mixture was adjusted to > 8.5 with concentrated ammonium hydroxide (5 drops) under vigorous stirring. The resulting phases were then separated, and the aqueous phase was extracted with dichloromethane (2 x 10 mL). The combined organic layers were washed with 0.25 M NaOH (15 mL), and brine (15 mL), then dried with sodium sulfate, filtered by gravity, and concentrated under reduced pressure. The crude residue was purified by flash chromatography (SiO_2_) with 10% methanol in chloroform as eluent to afford nornarwedine (8.3 mg, 42%). Spectroscopic data (GCMS, ^1^H-NMR and ^13^C-NMR) matched those reported in the literature^61,62^.

### Sample preparation for metabolite analysis

Metabolites were extracted from the crushed snap-frozen samples using a methanol-water extraction solvent (80:2, v/v) at a consistent ratio of 20 µL/mg of the sample. Before extraction, an internal standard of papaverine was added at a concentration of 2 ppm. The samples were then placed on a shaker at 200 rpm overnight, followed by heating at 65°C for 20 minutes with the lids closed. After centrifugation at 10,000 rpm for 10 minutes, the supernatants were collected and dried using a rotary evaporator. The dried extracts were reconstituted in the mobile phase, consisting of Milli-Q water and methanol (both containing 0.1% formic acid) at a 90:10 ratio by concentrating 20 times of initial extraction volume. The reconstituted samples were vortexed, filtered through a 0.2 µm PTFE filter, and subsequently analyzed by HPLC-MS/MS.

### Metabolite screening in *Leucojum aestivum* leaf

*Leucojum aestivum* bulbs purchased from The Seed Company (Canada) were grown indoor in the laboratory plant room. Untargeted metabolite analysis of *L. aestivum* leaves was performed by GC-MS, while targeted metabolite analysis was performed by LC-MS/MS. Cryopreserved macerated leaves (100 mg) were extracted with 2000 μL of solvent (80:20 methanol:water), heated at 60°C for 20 minutes, followed by 20 minutes of vortexing. The extract was centrifuged, and the supernatant was collected in another tube. For the remaining biomass, the non-polar solvent hexane was used for further extraction with an equal volume as before. The supernatant of the extracts collected in new tubes was evaporated under vacuum in a rotary evaporator for the polar fraction and under a stream of nitrogen for the non-polar fraction. The dried sample was reconstituted with mobile phase for LC-MS/MS and in 100% methanol for GCMS analysis ^63^.

### Instrumentation and chromatographic conditions

#### LC-MS/MS

High-performance liquid chromatography (HPLC) coupled with tandem mass spectrometry (MS/MS) (Agilent Technologies, Santa Clara, California, USA) system equipped with an Agilent Jet Stream ionization source, a binary pump, an autosampler and a column compartment were used for the analysis. Compounds separation was achieved using a Kinetex EVO C18 column (150 × 4.6 mm, 5 μm, 100 Å; Phenomenex, Torrance, USA). Five microliters of each sample were injected onto the column that was set at 30°C. A gradient method made of (A) formic acid 0.1% v/v in water and (B) formic acid 0.1% v/v in methanol with a flow rate of 0.4 mL/min was used to achieve chromatographic separation. The HPLC elution program is described as follows: 0 min, 10% B; 10.0 min, 10% B; 20.0 min, 100% B; 25.0 min, 100% B; 26.0 min, 10% B. The total run time was 30 min per sample to allow the reconditioning of the column prior to the next injection. The parameters used in the MS/MS source were set as follows: gas flow rate 10 L·min^-1^, gas temperature 300°C, nebulizer 45 psi, sheath gas flow 11 L·min^-1^, sheath gas temperature 300°C, capillary voltage 4000 V in ESI^+^ and 3500 V in ESI^-^ and nozzle voltage 500 V. Agilent MassHunter Data Acquisition (version 1.2) software was used to control the HPLC-MS/MS. MassHunter Qualitative Analysis (version 10.0) and MassHunter Quantitative QQQ Analysis (version 10.0) softwares were used for data processing. Compound identifications were made using authentic reference standards or by using collision-induced dissociation (CID) mass spectra published in literature ^64–66^ in the case where reference standards were not available. MRM transitions and instrument parameters used for identifications in *in vivo* experiments are included in Table S4. Instrument parameters used in MRM mode for metabolite screening in *L. aestivum* leaves and in SIM mode for in-vitro assays are presented in Tables S5 and S6 respectively. Instrument parameters used in Product Ion mode to obtain the full CID mass spectra of signals detected in SIM mode are presented in Table S7.

#### GC-MS

For GC-MS analysis, samples were injected into the GC-MS (Agilent Technologies 6890N GC coupled with 5973N inert MSD) in electron ionization mode at 70 eV. The temperature ramp used is described as follows: temperature was set at 100°C for 2 min, followed by 100–180°C at 15°C·min^−1^, 180–300°C at 5°C·min^−1^, and a 10 min hold at 300°C. Injector and detector temperatures were set at 250°C and 280°C, respectively, and the flow rate of carrier gas (He) was 1 mL·min^−1^. A split ratio of 1:10 was applied, and the injection volume was 1 μL.

### Statistical analysis

All statistical analyses were performed using GraphPad Prism 8 (GraphPad Software, San Diego, CA). Data are presented as mean ± standard deviation (SD) from at least three independent biological replicates unless otherwise stated. Comparisons between groups were evaluated using two-way analysis of variance (ANOVA), followed by Tukey’s multiple comparisons test to determine statistical significance between group means. A *p*-value of less than 0.05 was considered statistically significant.

## Acknowledgement

We thank Priya Gatti and Melodie B. Plourde for helping with confocal imaging and Arthur Renault for assistance with experiments during his research internship. We also thank Nuwan Sameera Liyanage for insightful discussion for substrate synthesis required for experiment and Karen Cristine Gonçalves dos Santos for support in editing the manuscript.

## 7 Authors Contribution

Basanta Lamichhane: Conceptualization, Methodology, Investigation, Formal analysis, Validation, Visualization, Writing- Original draft, Reviewing and Editing; Archana Niraula: Methodology, Investigation, Resources, and Editing; Natacha Merindol: Conceptualization, Methodology, Formal analysis, Project administration, Resources, Reviewing and Editing; Sarah-Eve Gélinas: Methodology, Resources, and Editing; Patrick Lagüe: Review of docking analysis; Simon Ricard: Substrate synthesis, Reviewing and Editing; Hugo Germain: Conceptualization, Methodology, Resources, Co-supervision, Reviewing and Editing; Isabel Desgagné-Penix: Conceptualization, Methodology, Resources, Funding acquisition, Supervision, Reviewing and Editing.

## References

1 Desgagné-Penix, I. Biosynthesis of alkaloids in Amaryllidaceae plants: A review. Phytochemistry Reviews 20, 409–431 (2021).

2 Cimmino, A., Masi, M., Evidente, M., Superchi, S. & Evidente, A. Amaryllidaceae alkaloids: Absolute configuration and biological activity. Chirality 29, 486–499 (2017).

3 Tallini, L. R. et al. Antitumoral activity of different Amaryllidaceae alkaloids: In vitro and in silico assays. Journal of Ethnopharmacology 329, 118154 (2024).

4 Sokhna, S. et al. Potential of several triazene derivatives against DENGUE viruses. Bioorganic & Medicinal Chemistry Letters 101, 129646 (2024).

5 Ka, S. et al. Amaryllidaceae alkaloid cherylline inhibits the replication of dengue and Zika viruses. Antimicrobial Agents and Chemotherapy 65, 10.1128/aac. 00398-00321 (2021).

6 Jayawardena, T. U., Merindol, N., Liyanage, N. S. & Desgagné-Penix, I. Unveiling Amaryllidaceae alkaloids: from biosynthesis to antiviral potential–a review. Natural Product Reports (2024).

7 Paiva, M., Nascimento, G., Damasceno, I., Santos, T. & Silveira, D. Pharmacological and toxicological effects of Amaryllidaceae. Brazilian Journal of Biology 83, e277092 (2023).

8 Wessjohann, L., Dippe, M., Tengg, M. & Gruber-Khadjawi, M. Methyltransferases in biocatalysis. Cascade biocatalysis: integrating stereoselective and environmentally friendly reactions, 393–426 (2014).

9 Mehta, N., Meng, Y., Zare, R., Kamenetsky-Goldstein, R. & Sattely, E. A developmental gradient reveals biosynthetic pathways to eukaryotic toxins in monocot geophytes. Cell 187, 5620–5637. e5610 (2024).

10 Liyanage, N. S. et al. Coclaurine N-methyltransferase-like enzymes drive the final biosynthetic reaction of the anti-Alzheimer’s drug galanthamine in Amaryllidaceae. Plant Physiology and Biochemistry, 110067 (2025).

11 Liscombe, D. K., Louie, G. V. & Noel, J. P. Architectures, mechanisms and molecular evolution of natural product methyltransferases. Natural product reports 29, 1238–1250 (2012).

12 Biastoff, S., Brandt, W. & Dräger, B. Putrescine N-methyltransferase–the start for alkaloids. Phytochemistry 70, 1708–1718 (2009).

13 Chen, W., Cheng, X., Zhou, Z., Liu, J. & Wang, H. Molecular cloning and characterization of a tropinone reductase from Dendrobium nobile Lindl. Molecular biology reports 40, 1145–1154 (2013).

14 Rasi, A., Sabokdast, M., Naghavi, M. R., Jariani, P. & Dedičová, B. Modulation of Tropane Alkaloids’ Biosynthesis and Gene Expression by Methyl Jasmonate in Datura stramonium L.: A Comparative Analysis of Scopolamine, Atropine, and Hyoscyamine Accumulation. Life 14, 618 (2024).

15 Levac, D. E. R. The γ-tocopherol-like family of N-methyltransferases: A taxonomically clustered gene family encoding enzymes responsible for N-methylation of monoterpene indole alkaloids. (2013).

16. Farzana, M. Phylogenetic and biochemical characterization of γ-tocopherol like-N-methyltransferase involved in the biosynthesis of biologically active monoterpenoid indole alkaloids. (2024).

17 Guo, J., Gao, D., Lian, J. & Qu, Y. De novo biosynthesis of antiarrhythmic alkaloid ajmaline. Nature Communications 15, 457 (2024).

18 Liscombe, D. K., Usera, A. R. & O’Connor, S. E. Homolog of tocopherol C methyltransferases catalyzes N methylation in anticancer alkaloid biosynthesis. Proceedings of the National Academy of Sciences 107, 18793–18798 (2010).

19 Levac, D., Cázares, P., Yu, F. & De Luca, V. A picrinine N-methyltransferase belongs to a new family of γ-tocopherol-like methyltransferases found in medicinal plants that make biologically active monoterpenoid indole alkaloids. Plant Physiology 170, 1935–1944 (2016).

20 Kato, M. et al. Caffeine biosynthesis in young leaves of Camellia sinensis: In vitro studies on N-methyltransferase activity involved in the conversion of xanthosine to caffeine. Physiologia plantarum 98, 629–636 (1996).

21 Kato, M. et al. Expression for caffeine biosynthesis and related enzymes in Camellia sinensis. Zeitschrift für Naturforschung C 65, 245–256 (2010).

22 Zhou, M.-z., et al. N-methyltransferases of caffeine biosynthetic pathway in plants. Journal of Agricultural and Food Chemistry 68, 15359–15372 (2020).

23 Ashihara, H., Sano, H. & Crozier, A. Caffeine and related purine alkaloids: biosynthesis, catabolism, function and genetic engineering. Phytochemistry 69, 841–856 (2008).

24 Choi, K.-B., Morishige, T., Shitan, N., Yazaki, K. & Sato, F. Molecular cloning and characterization of coclaurinen-methyltransferase from cultured cells of Coptis japonica. Journal of Biological Chemistry 277, 830–835 (2002).

25 Morris, J. S. & Facchini, P. J. Isolation and characterization of reticuline N-methyltransferase involved in biosynthesis of the aporphine alkaloid magnoflorine in opium poppy. Journal of Biological Chemistry 291, 23416–23427 (2016).

26 Liscombe, D. K. & Facchini, P. J. Molecular cloning and characterization of tetrahydroprotoberberine cis-N-methyltransferase, an enzyme involved in alkaloid biosynthesis in opium poppy. Journal of Biological Chemistry 282, 14741–14751 (2007).

27 Morris, J. S. & Facchini, P. J. Molecular Origins of Functional Diversity in Benzylisoquinoline Alkaloid Methyltransferases. Frontiers in Plant Science 10 (2019).

28 Bennett, M. R. et al. Structure and Biocatalytic Scope of Coclaurine N-Methyltransferase. Angew Chem Int Ed Engl 57, 10600–10604 (2018).

29 Torres, M. A. et al. Structural and functional studies of pavine N-methyltransferase from Thalictrum flavum reveal novel insights into substrate recognition and catalytic mechanism. Journal of Biological Chemistry 291, 23403–23415 (2016).

30 Lamichhane, B. et al. Elucidating the enzyme network driving Amaryllidaceae alkaloids biosynthesis in Leucojum aestivum. Plant Biotechnology Journal (2025).

31 Ptak, A. et al. Carbohydrates stimulated Amaryllidaceae alkaloids biosynthesis in Leucojum aestivum L. plants cultured in RITA® bioreactor. PeerJ 8, e8688 (2020).

32 Szlávik, L. et al. Alkaloids from Leucojum vernum and antiretroviral activity of Amaryllidaceae alkaloids. Planta medica 70, 871–873 (2004).

33 Nelson, B. K., Cai, X. & Nebenführ, A. A multicolored set of in vivo organelle markers for co-localization studies in Arabidopsis and other plants. The Plant Journal 51, 1126–1136 (2007).

34 Kozbial, P. Z. & Mushegian, A. R. Natural history of S-adenosylmethionine-binding proteins. BMC structural biology 5, 1–26 (2005).

35 Wlodarski, T. et al. Comprehensive structural and substrate specificity classification of the Saccharomyces cerevisiae methyltransferome. PloS one 6, e23168 (2011).

36 Koudounas, K. et al. Tonoplast and peroxisome targeting of γ-tocopherol N-methyltransferase homologs involved in the synthesis of monoterpene indole alkaloids. Plant and Cell Physiology 63, 200–216 (2022).

37 Desgagné-Penix, I. & Facchini, P. J. Systematic silencing of benzylisoquinoline alkaloid biosynthetic genes reveals the major route to papaverine in opium poppy. The Plant Journal 72, 331–344 (2012).

38 Schluckebier, G., O’Gara, M., Saenger, W. & Cheng, X. Vol. 247 16-20 (Academic Press, 1995).

39 Martin, J. L. & McMillan, F. M. SAM (dependent) I AM: the S-adenosylmethionine-dependent methyltransferase fold. Current opinion in structural biology 12, 783–793 (2002).

40 Lee, S. et al. O-and N-Methyltransferases in benzylisoquinoline alkaloid producing plants. Genes & Genomics 46, 367–378 (2024).

41 Berkov, S., Bastida, J., Viladomat, F. & Codina, C. Development and validation of a GC–MS method for rapid determination of galanthamine in Leucojum aestivum and Narcissus spp.: A metabolomic approach. Talanta 83, 1455–1465 (2011).

42 Karimzadegan, V. et al. Characterization of cinnamate 4-hydroxylase (CYP73A) and p-coumaroyl 3′-hydroxylase (CYP98A) from Leucojum aestivum, a source of Amaryllidaceae alkaloids. Plant Physiology and Biochemistry 210, 108612 (2024).

43 Hagel, J. M. & Facchini, P. J. Subcellular localization of sanguinarine biosynthetic enzymes in cultured opium poppy cells. In Vitro Cellular & Developmental Biology-Plant 48, 233–240 (2012).

44 Kodama, Y., Shinya, T. & Sano, H. Dimerization of N-methyltransferases involved in caffeine biosynthesis. Biochimie 90, 547–551 (2008).

45 Guirimand, G. et al. The subcellular organization of strictosidine biosynthesis in Catharanthus roseus epidermis highlights several trans-tonoplast translocations of intermediate metabolites. The FEBS journal 278, 749–763 (2011).

46 Hugueney, P. et al. A novel cation-dependent O-methyltransferase involved in anthocyanin methylation in grapevine. Plant physiology 150, 2057–2070 (2009).

47 Guirimand, G. et al. Spatial organization of the vindoline biosynthetic pathway in Catharanthus roseus. Journal of plant physiology 168, 549–557 (2011).

48 Zbierzak, A. M. et al. Intersection of the tocopherol and plastoquinol metabolic pathways at the plastoglobule. Biochemical Journal 425, 389–399 (2010).

49 Muñoz, P. & Munné-Bosch, S. Vitamin E in plants: biosynthesis, transport, and function. Trends in plant science 24, 1040–1051 (2019).

50 Vitale, A., Ceriotti, A. & Denecke, J. The role of the endoplasmic reticulum in protein synthesis, modification and intracellular transport. Journal of Experimental Botany 44, 1417–1444 (1993).

51 Held, M. A. et al. CGR3: a Golgi-localized protein influencing homogalacturonan methylesterification. Molecular plant 4, 832–844 (2011).

52 Temple, H. et al. Golgi-localized putative S-adenosyl methionine transporters required for plant cell wall polysaccharide methylation. Nature Plants 8, 656–669 (2022).

53 Majhi, B. B., Gélinas, S.-E., Mérindol, N., Ricard, S. & Desgagné-Penix, I. Characterization of norbelladine synthase and noroxomaritidine/norcraugsodine reductase reveals a novel catalytic route for the biosynthesis of Amaryllidaceae alkaloids including the Alzheimer’s drug galanthamine. Frontiers in Plant Science 14, 1231809 (2023).

54 Eichhorn, J., Takada, T., Kita, Y. & Zenk, M. H. Biosynthesis of the Amaryllidaceae alkaloid galanthamine. Phytochemistry 49, 1037–1047 (1998).

55 Tamura, K., Stecher, G. & Kumar, S. MEGA11: molecular evolutionary genetics analysis version 11. Molecular biology and evolution 38, 3022–3027 (2021).

56 Jones, D. T., Taylor, W. R. & Thornton, J. M. The rapid generation of mutation data matrices from protein sequences. Bioinformatics 8, 275–282 (1992).

57 Letunic, I. & Bork, P. Interactive Tree Of Life (iTOL) v5: an online tool for phylogenetic tree display and annotation. Nucleic acids research 49, W293–W296 (2021).

58 Abramson, J. et al. Accurate structure prediction of biomolecular interactions with AlphaFold 3. Nature 630, 493–500 (2024).

59 Irwin, J. J. et al. ZINC20-A Free Ultralarge-Scale Chemical Database for Ligand Discovery. J Chem Inf Model 60, 6065–6073 (2020).

60 Adasme, M. F. et al. PLIP 2021: expanding the scope of the protein–ligand interaction profiler to DNA and RNA. Nucleic Acids Research 49, W530–W534 (2021).

61 Kogure, N., Katsuta, N., Kitajima, M. & Takayama, H. Two new alkaloids from Crinum asiaticum var. sinicum. Chemical and Pharmaceutical Bulletin 59, 1545–1548 (2011).

62 Hirnschall, M., Froehlich, J., Jordis, U. & Mereiter, K. Seco-isopowellaminone: A new heterocycle from nornarwedine. Journal of heterocyclic chemistry 39, 1265–1270 (2002).

63 Baeshen, N. A. et al. GC-MS analysis of bioactive compounds extracted from plant Rhazya stricta using various solvents. Plants 12, 960 (2023).

64. Mehta, N., Meng, Y., Zare, R., Kamenetsky-Goldstein, R. & Sattely, E. A developmental gradient reveals biosynthetic pathways to eukaryotic toxins in monocot geophytes. bioRxiv (2023).

65 Kilgore, M. B., Augustin, M. M., May, G. D., Crow, J. A. & Kutchan, T. M. CYP96T1 of Narcissus sp. aff. Pseudonarcissus catalyzes formation of the Para-Para’CC phenol couple in the Amaryllidaceae alkaloids. Frontiers in plant science 7, 225 (2016).

66 Kilgore, M. B., Holland, C. K., Jez, J. M. & Kutchan, T. M. Identification of a noroxomaritidine reductase with Amaryllidaceae alkaloid biosynthesis related activities. Journal of Biological Chemistry 291, 16740–16752 (2016).

